# Standardizing the mouse MCAO Stroke Model: Distinguishing Internal Carotid vs Pterygopalatine Artery Occlusion

**DOI:** 10.1101/2025.10.20.683461

**Authors:** Ayman Isbatan, Samuel M Lee, Frederick C. Damen, Sunny Chen, Michelle Malson, Xiaoping Du, Richard D. Minshall, Kejia Cai, Jiwang Chen

## Abstract

**Background:** The intraluminal middle cerebral artery occlusion (MCAO) model is widely used in preclinical stroke research, but outcomes remain inconsistent because of variable occlusion times and inadvertent filament placement into the pterygopalatine artery (PPA).

**Methods:** We applied a modified version of the Koizumi method to directly visualize the internal carotid artery (ICA)–PPA bifurcation and systematically compare targeted ICA versus PPA occlusion. We combined multiple complementary methods - survival studies, neurological scoring, 2,3,5-Triphenyltetrazolium chloride (TTC) infarct staining, Evans blue blood brain barrier (BBB) permeability assays, laser speckle cerebral blood flow imaging, small-animal magnetic resonance imaging (MRI), and evaluation of inflammatory gene expression through qPCR - to rigorously characterize injury patterns, survival, and inflammatory responses.

**Results:** ICA occlusion approximately caused a rapid 70% reduction in cerebral blood flow, progressive infarction (15 min: smaller infarct with approximately 50% 7-day survival; 30 min: moderate infarct but >50% mortality), and cortical edema. In contrast, PPA occlusion produced only a gradual 44% decline in blood flow, small infarcts at 60 min) and preserved survival (80% at 7 days). Finally, magnetic resonance imaging in 15-minute ICA occluded mice identified the development of cortical edema which was not observed in sham, 15-minute PPA, or 60-minute PPA mice.

**Conclusion:** These findings identify 15-minute ICA occlusion as the most practical balance between reproducible infarction and survival for long-term studies, while 30 minutes is best reserved for short-term infarct induction when survival is less critical. By resolving two critical sources of MCAO variability, occlusion duration and vascular targeting, this study provides a standardized protocol and evidence-based occlusion benchmarks that will improve reproducibility in preclinical stroke studies

## Introduction

Stroke remains the second leading cause of global mortality and long-term disability, with ischemic stroke accounting for approximately 80% of stroke related cases ^1–3^. In the United States alone, stroke-related costs are projected to reach $1.49 trillion by 2050, underscoring the urgent need for improved therapies ^4^. Since occlusion of the middle cerebral artery accounts for half of ischemic strokes ^5^, preclinical middle cerebral artery occlusion (MCAO) research remains a central priority.

The intraluminal MCAO model remains the most widely used preclinical model in the field of stroke due to its relative technical ease and ability to produce a cerebral infarction in the lateral cerebral hemisphere ^6,7^. Originally developed by Koizumi in 1986 ^8^, a silicone-coated filament is inserted into the common carotid artery (CCA) and advanced into the internal carotid artery (ICA) to occlude the origin of the MCA. However, experience in our lab suggests that the filament can deviate into the pterygopalatine artery (PPA) which branches from the CCA and provides blood flow to the nasal and oral cavity ^9^. This anatomical challenge has been largely overlooked in prior studies and may explain the variability observed between labs using the intraluminal MCAO model.

Two major sources of variability undermine the intraluminal MCAO model’s reproducibility. First, occlusion times vary widely across studies, typically ranging from 30 minutes to 2 hours ^6,7,10^. In a systematic review of 129 rodent MCAO studies from 1989-2025, the most common occlusion times used were 30 min ^11–13^ (13.2% studies), 60 min ^14–16^ (18.6%), 90 min ^17–19^ (18.6%), and 120^20–22^ (24.0%) (supplemental table 1). Secondly, the unintended occlusion of the PPA produces a milder level of ischemic injury since it does not directly perfuse the cerebrum and instead supplies blood to the retina through the ophthalmic artery ^23^. Despite these limitations, no prior study has directly compared ICA versus PPA occlusion side-by-side to define their distinct contributions to infarct variability.

In this study, we addressed this critical gap by systematically comparing ICA versus PPA occlusion using a modified Koizumi protocol that improves anatomical visualization of the ICA–PPA bifurcation. We combined multiple complementary methods — including filament placement, survival studies, neurological scoring, TTC infarct staining, Evans blue BBB permeability assays, laser speckle cerebral blood flow imaging, small-animal magnetic resonance imaging (MRI), and evaluation of inflammatory gene expression through qPCR — to rigorously characterize injury patterns, survival, and inflammatory responses. By integrating these approaches, we aimed to resolve the variability of the MCAO model and provide reproducible benchmarks for stroke investigators. By resolving these sources of variability, our protocol strengthens the reliability of preclinical stroke models, which is essential for advancing therapies into clinical testing.

## Materials and Methods

### Mice

12 week old male C57BL/6 male mice weighing approximately 20 grams were used for this study and approved for use by the Institutional Animal Care and Use Committee of the University of Illinois at Chicago. All experiments were performed in accordance with the guidelines and regulations of the University of Illinois at Chicago.

### Intraluminal Middle Cerebral Artery Occlusion (MCAO) Model

Anesthetized intubated animals (isoflurane 3% induction, 1.5-2% maintenance) were placed on a heated surgical table (37°C) and connected to a Minivent-ventilator (Model 845 – [Harvard Apparatus, Massachusetts, USA). Tidal volume (Vt; mL) was maintained at 0.120 mL, whereas respiratory rate was maintained at 148 breaths per minute.

A midline cervical incision was performed to expose the left common carotid artery (LCCA), which was isolated from surrounding vagal nerve tissue using suture tying forceps. The left external carotid artery (LECA) and left internal carotid artery (LICA) were then identified. To assist in the visualization of the LICA and left pterygopalatine artery (LPPA), the hypoglossal (HG) nerve - which runs before the branching point of the LICA and LPPA - was identified. The HG nerve was mobilized by gently separating the surrounding tissue, thereby improving visibility of the LICA and LPPA.

The LECA was then temporarily occluded with a reversible 5-0 silk ligature, while a loose 5-0 silk ligature was placed around the LICA. A small arteriotomy was then made in the LCCA using micro-scissors, through which a silicone rubber-coated monofilament (diameter: 0.19 mm for 15-20 g mice, Doccol Corp #6021910PK5Re) was inserted. The filament was gently advanced into either the LICA or left pterygopalatine artery (LPPA) under direct visualization of the bifurcation (figure 1a). Upon encountering resistance at the MCA origin (typically at 9-11 mm insertion depth from carotid bifurcation), the filament was secured using the LCCA ligature.

**Figure 1:**
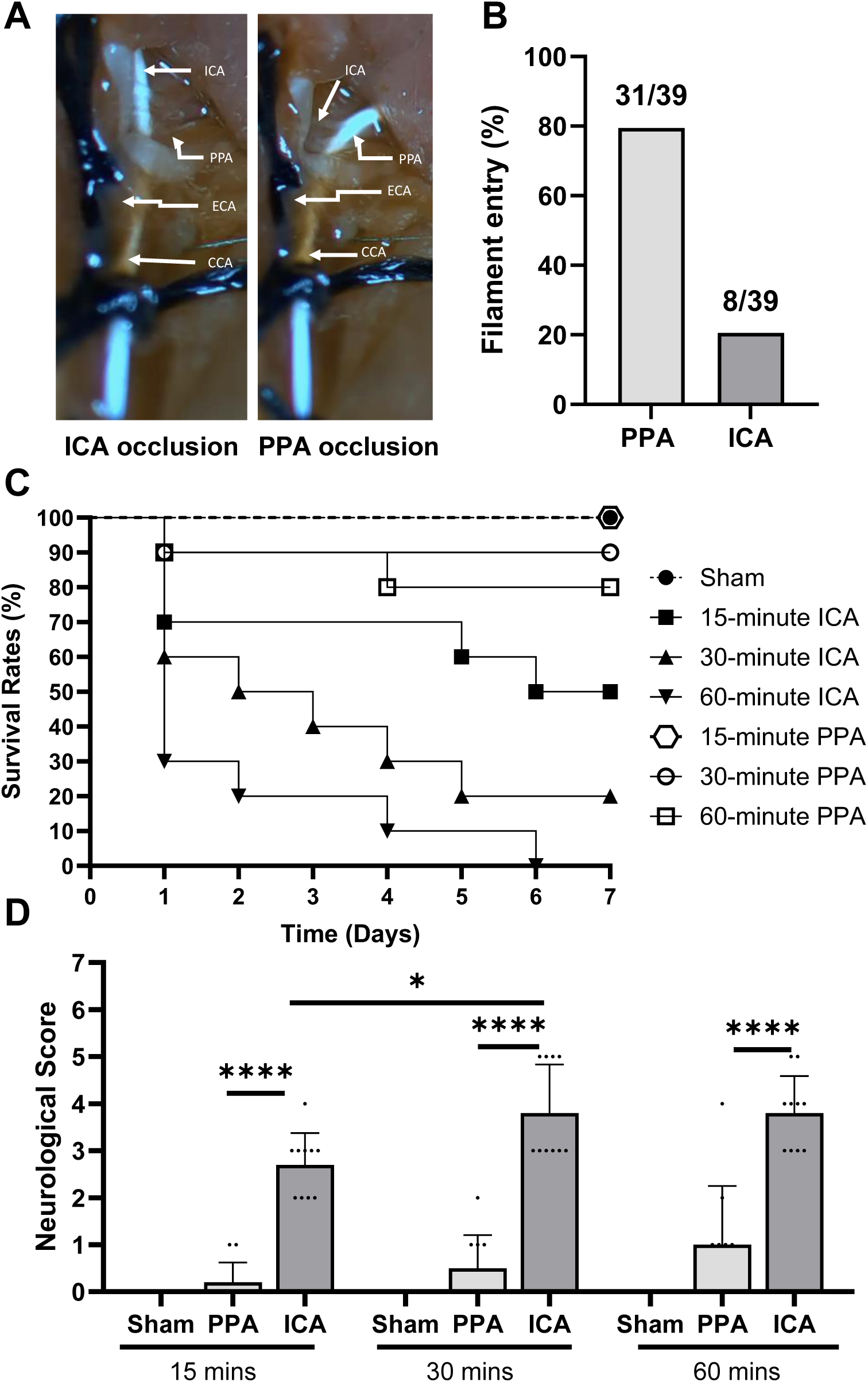
Acute occlusion of the internal carotid artery (ICA) causes the rapid and progressive decline in survival rates and neurological deficits in mice. (A) Representative images ICA and pterygopalatine artery (PPA) occluded with a filament. (B) Chances of filament entry in either ICA or PPA. (C) Kaplan–Meier survival curves comparing internal carotid artery (ICA) versus PPA occlusion for 15, 30, and 60 minutes, followed for 7 days. (D) Neurological scores (Bederson scale) 24 h post-surgery. ICA occlusion caused progressive mortality and severe neurological deficits, whereas PPA occlusion preserved survival and produced only mild deficits. Sample size for neurological scoring: n = 10. Data shown as mean ± SD. * - P < 0.05, **** - P < 0.0001.

Occlusion was maintained for 15, 30, or 60 minutes. After the designated occlusion period, the filament was withdrawn to allow reperfusion, and the LCCA was permanently ligated with 5-0 silk suture. The surgical site was closed in layers, and animals were recovered under a heat lamp with supplemental medical-grade oxygen. Detailed surgical protocols are provided in the Supplementary Materials.

### Evaluation of neurological deficit

Neurological evaluation was performed at 24 h after reperfusion using the Bederson scoring system ^24^. Scores ranged from 0 to 5: 0 = normal motor function; 1 = mild circling (<50% attempts to rotate to the contralateral side); 2 = consistent circling (>50% attempts to rotate to the contralateral side); 3 = strong circling with nose-to-tail rotation lasting >1–2 s; 4 = severe rotation, barreling, or loss of righting reflex; 5 = comatose or moribund.

### TTC Staining / Evaluation of infarct size

After 24 hours mice were euthanized, the brains were extracted and placed in an acrylic mouse brain matrix (Ted Pella Inc, California, USA) and cut into coronal sections (1mm thick). The sections were stained in 0.9% NaCl-PBS with 2% 2,3,5-triphenyl tetrazolium chloride (TTC) (TCI America, Oregon, USA) for 30 minutes at room temperature and fixed overnight in 10% neutral buffered formalin solution (Sigma-Aldrich, Massachusetts, USA).

In viable tissue, TTC is enzymatically reduced to form red 1,3,5-triphenylformazan, while necrotic areas remain white. Infarct volume was quantified using ImageJ (version 1.54p 17), expressed as infarction area/total area ^25^. Analyses were performed with blinding to experimental groups.

### Evaluation of blood-brain barrier (BBB) integrity

Evans Blue (EB, 2% in saline, 4 ml/kg) was administered via retro-orbital injection 24 h after MCAO onset. After 1 h, mice were anesthetized with ketamine (80–100 mg/kg) and xylazine (10 mg/kg). A thoracotomy was performed, and the inferior vena cava was cut at the abdominal region. The mouse was perfused through the left ventricle with saline using a syringe pump at 3 ml/min at 110 mmHg (Graseby 3400 syringe pump; Graseby, Watford, UK) until colorless fluid appeared from the vena cava.

After decapitation, brains were bisected into hemispheres, weighed, and placed in 50% trichloroacetic acid. After homogenization and centrifugation, dye extracts were diluted with ethanol (1:3), and fluorescence was measured (excitation 620 nm, emission 680 nm). EB concentration was calculated using a standard curve (100–500 ng/ml) and expressed as ng/g of tissue.^26^

### Laser Speckle Contrast Real Time Imaging

A RFLSI ZW Laser Speckle Contrast system (RWD, Shenzhen, China) was used to measure CBF changes in real time before and after MCAO. Following anesthesia and ventilation as described above, mice were positioned prone. Hair over the cortical region was removed, the scalp disinfected, and an incision made to expose the skull. The skull surface was pre-wetted with saline, and the laser head was positioned 10–15 cm above the left hemisphere.

Baseline CBF was recorded for 3–5 min. Mice were then repositioned supine for MCAO, followed by return to prone position for ≥2 h of real-time imaging. Blood flow changes were quantified relative to baseline using RFLSI software ^27^. Analyses were blinded to experimental groups.

### Magnetic Resonance Imaging (MRI)

Non-invasive real-time MRI was performed in sham, ICA (15 min) and PPA (15 and 60 min) occluded mice at 24 hours post-occlusion. Due to the high rates of mortality, 60 minutes of ICA occlusions were not performed for MRI. Mice were anesthetized with 1–2% isoflurane in O₂, and body temperature was maintained at 37 °C. Relative cerebral blood flow (rCBF) was determined using a FAIR (Flow-Sensitive Alternating Inversion Recovery) ASL (Arterial Spin Labeling) preparation ^28^ with a FID-SSFP (free induction decay steady state free precession) readout, collected on a Varian 9.4T MRI small animal scanner and a 39-mm volume coil. The FAIR preparation used a hyperbolic secant adiabatic inversion pulse, duration 4 ms, with no slice selection gradient (tagged blood) and slice selection of twice the width of the image slice (non tagged blood), and an inversion time of 1.5 seconds with a T1-delay time of 6 seconds for full T1 recovery. The FID-SSFP used a centrically encoded readout with variable flip angles, TR/TE 2.53/1.27 ms, FA 1 ms Gauss pulse, 128×128 matrix, FOV 18×18 mm^2^, and slice thickness 1 mm. Similar to a previous study ^29^, the variable flip angles were empirically chosen to start the readout in the steady state, to maximize the ASL contrast and minimize blurring in the phase direction. Due to signal to noise (SNR) issues 200 averages were used. The rCBF was calculated by the difference in the untagged and tagged image divided by the water content, a proton-density weighted image acquired using a fast spin echo sequence with TR/TE 6000/5.3 ms and ETL 8. Besides the quantification of rCBF, tissue edema due to stroke was also determined using the ratio of T2-weighted MRI signals between the stroke (left) and contralateral (right) hemispheres. All quantitative images analyses were conducted with blinding to experimental groups.

### RNA isolation / TaqMan Real-Time PCR

Left lobes were isolated from experimental groups, flash frozen, and initially processed with a tissue pulverizer. Approximately 50-100 mg of processed tissue was suspended in TRIzol Reagent (Invitrogen, Carlsbad, USA) and homogenized with a mechanical tissue homogenizer (Fisher Scientific, Massachusetts, USA). Following RNA precipitation with molecular grade isopropanol, and subsequent washes with 70% ethanol, the dried RNA pellet was resuspended in diethyl pyrocarbonate water. RNA was further purified using Promega’s RQ1 RNase-Free DNase kit (Promega, Madison, USA) and then reverse transcribed using a RevertAid RT kit (Thermo Fisher Scientific, Skokie, USA). Samples were then run at 100 ng per well in 384 PCR plates with TaqMan probes (interleukin 6 (IL6), tumor growth factor beta (TGF-β)) (Thermo Fisher Scientific) and TaqMan Fast Advanced Mastermix (Applied Biosystems, Foster City, USA) using a Life Technologies ABI ViiA7 (Life Technologies, Carlsbad, USA).

### Statistical analysis

All graphs and data analyses were generated using Graphpad Prism software, version 10.1.2 for Windows (GraphPad Software, Massachusetts, USA, www.graphpad.com). Survival rates (24 h) and MRI data were analyzed using t-tests with Bonferroni correction. qPCR results were analyzed using one-way ANOVA with Dunnett post hoc test. Infarct size and BBB permeability were analyzed using two-way ANOVA with Tukey HSD post hoc test. Laser speckle blood flow changes were analyzed using two-way ANOVA with mixed-effect modeling. Data is presented as mean ± SD.

## Results

### Filament entry is more likely to enter the PPA than the ICA

To determine whether the filament would naturally enter the PPA or ICA, the unmodified version of the Koizumi method was performed to assess filament placement after the first attempt (figure 1A). Of the 39 mice, only 20.5% (8 mice) had the filament successfully entered the ICA whereas 79.5% (31 mice) had the filament enter the PPA (figure 1B).

### ICA occlusion time determines survivability and neurological deficits

ICA occlusion resulted in poor survivability and severe neurological deficits (figure 1C, 1D). For 15-minute ICA occlusions, 70% of mice survived after 24 hours which progressively declined to 50% after 7 days (figure 1C). The decline in survival was more apparent the longer the ICA occlusion times: 30 minutes (day 1: 60% survival; day 7: 20%) and 60 minutes (day 1: 30% survival; day 7: 0%). In comparison, PPA-occluded mice had higher survival rates: 15 minutes of PPA occlusion did not cause mortality, while 90% of mice survived for 30-minute occlusion after 7 days. Even after 1 hour of PPA occlusion, most of the mice survived (day 1: 90% survival; day 7: 80%).

These mice were also examined for signs of neurological deficiency 24 hours after occlusion (figure 1D). Two-way ANOVA showed that the location of the occlusion (ICA versus PPA, P < 0.0001) and the duration of ischemia were both independent variables which influenced neurological deficiency. When time matched, post-hoc analyses showed that ICA caused more severe neurological deficits compared to PPA for 15 min (PPA: 0.20 ± 0.42 vs ICA: 2.70 ± 0.68), 30 min (PPA: 0.50 ± 0.71 vs ICA: 3.80 ± 1.03), and 60 min (PPA: 1.00 ± 1.25 vs ICA: 3.80 ± 0.79) occlusions (P < 0.0001 for all comparisons). Post-hoc analyses also revealed that the decline in neurological function from ICA, but not PPA, occlusion was exacerbated after if cerebral ischemia was increased beyond 15 minutes (ICA-15 minutes vs ICA-30 minutes, P <0.0155). Occluding the ICA for 60 minutes did not lead to further neurological decline.

### Infarct size and Blood Brain Barrier (BBB) disruption depends on occlusion site and duration

The severe decline in neurological function and survivability is supported by the differences observed between infarct size in ICA versus PPA occluded brains (figure 2). As expected, ICA occlusion caused cerebral infarctions which were visualized as early as 15 minutes of ICA occlusion (figure 2A). Two-way ANOVA showed that the increase in cerebral infarction was influenced by filament placement (ICA versus PPA) and the length of ischemia (interactive effect: 0.0017) (figure 2B). Post-hoc analysis showed that the progressive increase in infarction size in ICA occluded brains was dependent on the duration of ischemia (ICA-15 minutes: 17.90 ± 9.01% vs ICA-30 minutes: 28.11 ± 7.27%, P < 0.0215; ICA-30 minutes vs ICA-60 minutes: 41.59 ± 6.92%, P < 0.0011). However, PPA occlusion failed to generate a cerebral infarction which could be detected by TTC staining after 15 minutes (0.34 ± 0.47%) or 30 minutes (0.71 ± 0.56%) of PPA occlusion. Although 60-minutes of PPA occlusions generated a cerebral infarction (10.355 ± 7.70%) this is significantly smaller compared to ICA-60-minute occluded mice when compared together (60 minutes: ICA vs PPA, P < 0.0001) (figure 2B).

**Figure 2.**
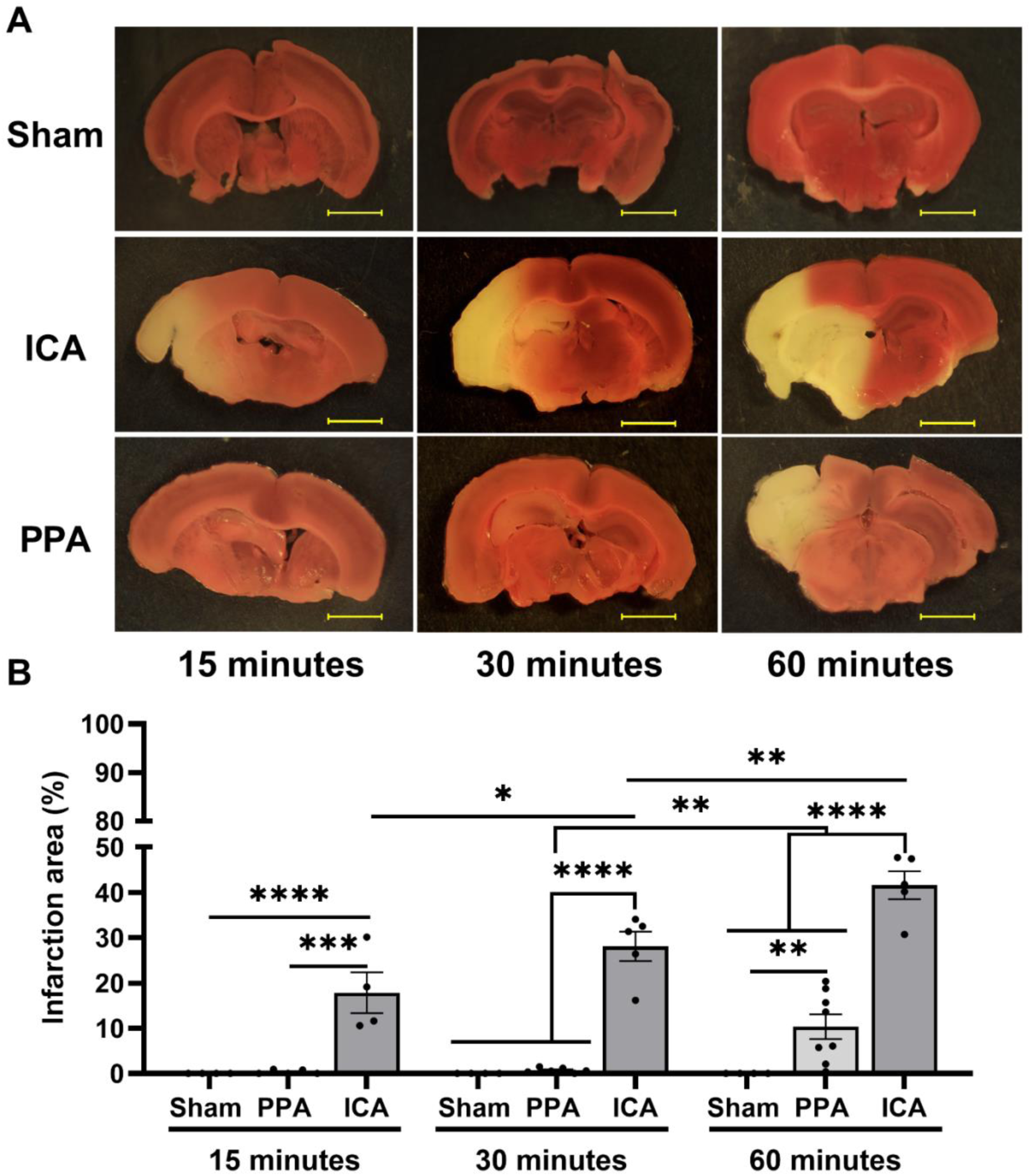
Infarct volume depends on occlusion site and ischemia duration. (A) Representative TTC-stained coronal sections from sham, ICA, and PPA occluded mice. (B) Quantification of infarct volume (% of hemisphere). ICA occlusion produced duration-dependent infarction (15, 30, 60 min), whereas PPA occlusion caused no infarct at 15–30 min and only small infarcts after 60 min. Yellow scale bar shown in images is 0.2 cm. Sample size: n =6–8 per group. Data shown as mean ± SD. *P < 0.05, **P < 0.001, ***P = 0.0001, ****P < 0.0001.

Assessment of BBB permeability of brains following ICA or PPA occlusion revealed similar trends since ICA, but not PPA, occlusion increased BBB permeability the longer the ICA occlusion (interactive effect: P = 0.002) (figure 3). Post-hoc analyses showed that BBB permeability significantly increased after 30 min (Sham: 0.095 ± 0.048 μg/g vs ICA:6.20 ± 0.69 μg/g) and 60 minutes of ICA occlusion (sham: 0.12 ± 0.064 μg/g vs ICA: 8.36 ± 0.42 μg/g, P < 0.0001) when compared to sham mice (figure 3B). Although PPA occlusion increased BBB permeability compared to sham mice, these values were significantly low. Although 30 minutes of PPA occlusion (Sham-30 minutes 0.095 ± 0.048 μg/g vs PPA-30 minutes: 4.14 ± 3.45 μg/g, P = 0.057) trended to have higher values of BBB permeability, statistical significance was only observed after 60 minutes (PPA-60 minute: 4.41 ± 3.45 μg/g, P = 0.0069).

**Figure 3.**
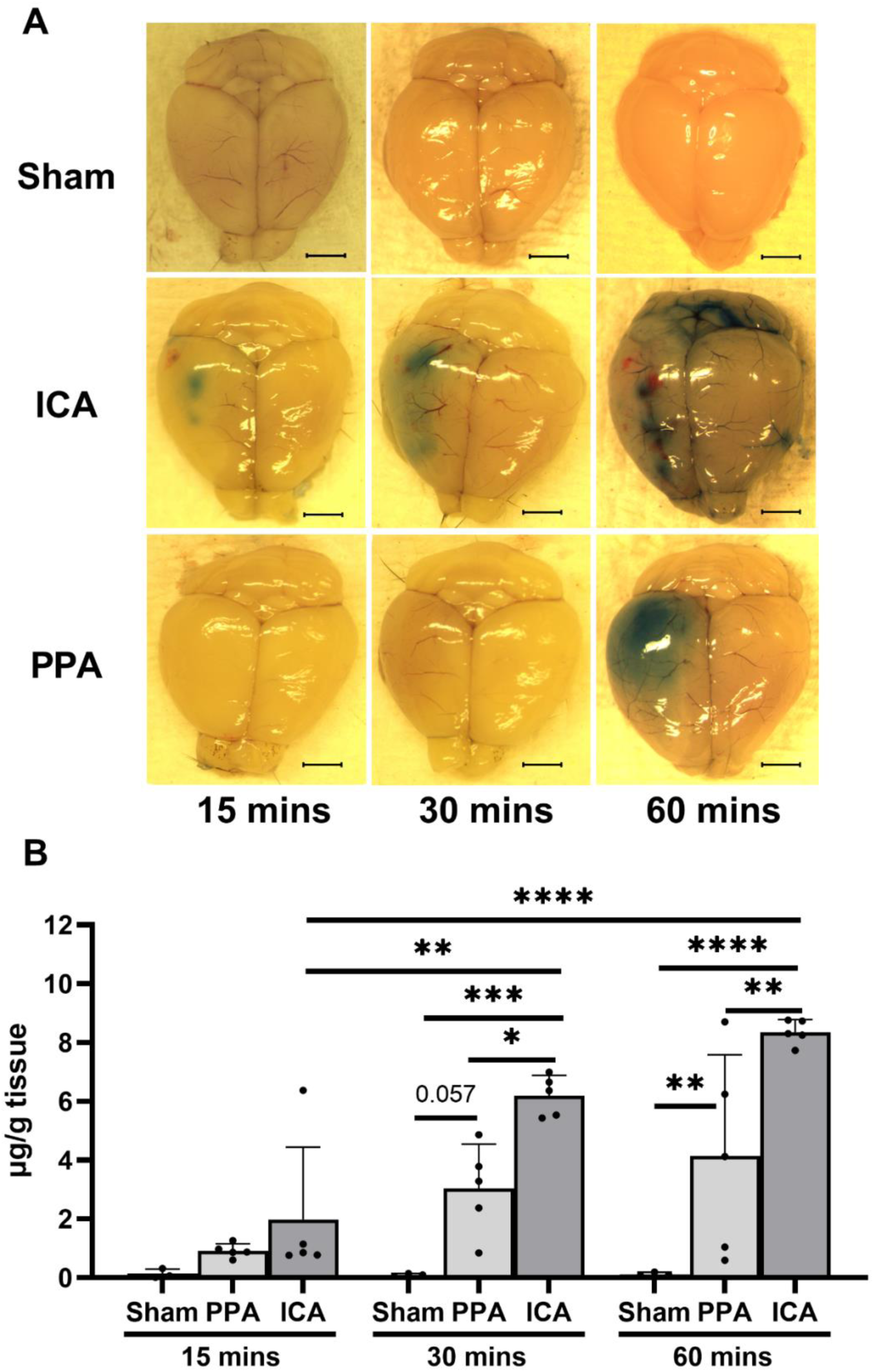
Blood–brain barrier (BBB) disruption is greater after ICA than PPA occlusion. (A) Representative Evans blue staining in sham, ICA, and PPA brains. (B) Quantified BBB permeability (µg/g). Prolonged ICA occlusion caused significant BBB leakage, while PPA occlusion showed only modest changes. Black scale bar shown in images is 0.2 cm. Sample size: n = 3–5 per group. Data shown as mean ± SD. *P < 0.05, **P < 0.001, ****P < 0.0001.

### Laser speckle imaging reveals rapid cerebral blood flow (CBF) loss after ICA occlusion

To evaluate whether there is a progressive decline in CBF from ICA and PPA occlusion, real time laser speckle imaging was performed (figure 4A). Two-way ANOVA with mixed effect modelling revealed that the method of occlusion (ICA vs PPA) and the duration of ischemia influenced the rate of CBF decline (interactive effect: 0.0001) (figure 4B). ICA occlusion caused a rapid decline in blood flow and remained consistent throughout the 2 hours of real time imaging (Sham blood flow baseline: 1.00 vs ICA 15 minutes: 0.23 ± 0.053 – 120 minutes: 0.22 ± 0.028, P < 0.05). PPA occlusion initially caused a moderate initial decline that progressed over time (15 minutes: 0.60 ± 0.032 - 120 minutes: 0.44 ± 0.13, P < 0.05). Despite its decline, PPA occlusion did not reach the same level of ischemic level as ICA (ICA-120 minutes: 0.22 ± 0.028 vs PPA-120 minutes: 0.44 ± 0.13, P = 0.079).

**Figure 4.**
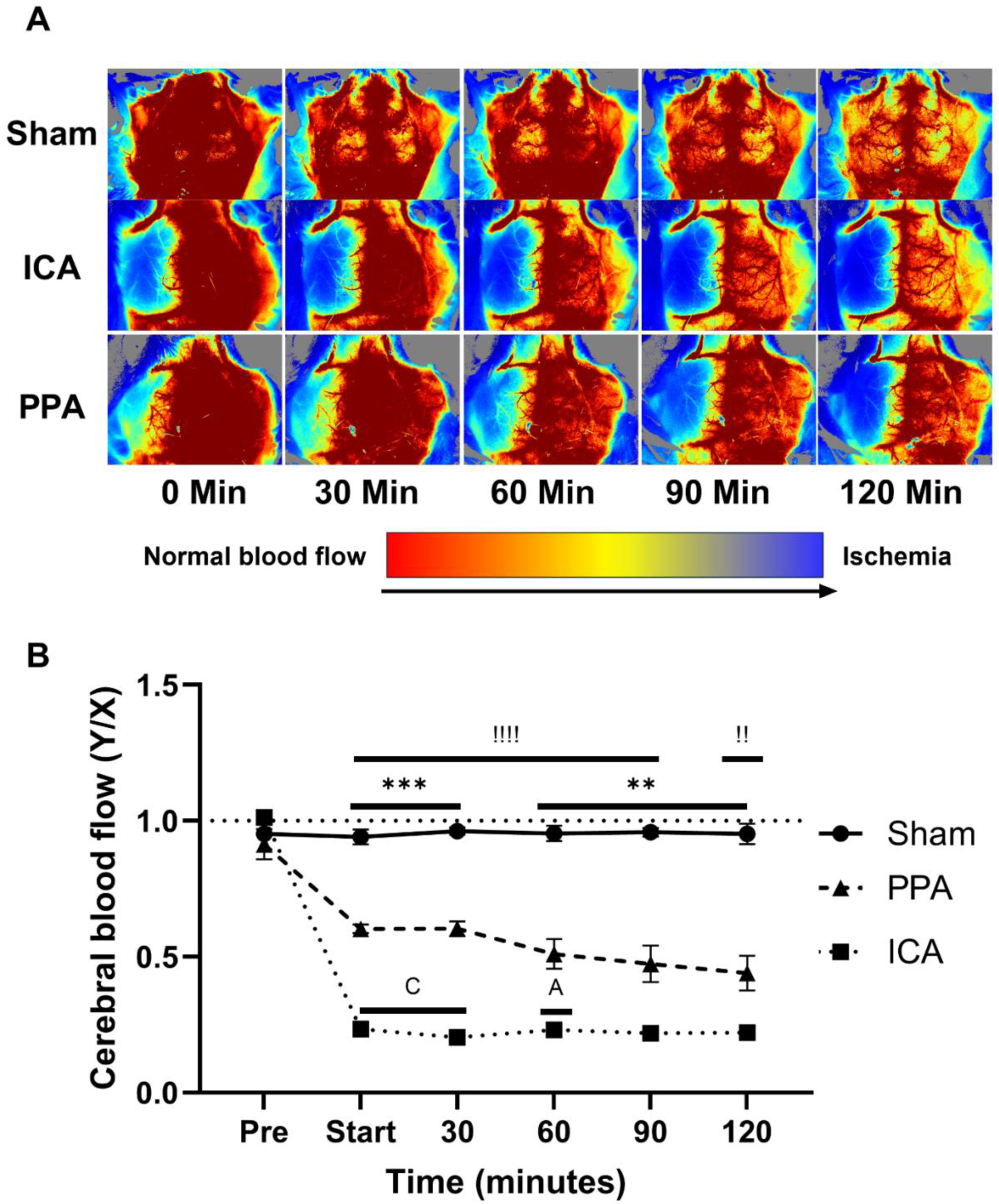
Cerebral blood flow (CBF) dynamics differ between ICA and PPA occlusion. (A) Representative laser speckle imaging maps showing blood flow changes during 120 min monitoring. Warm colors = preserved flow; cool colors = ischemia. (B) Quantified CBF over time. ICA occlusion caused a rapid and sustained CBF drop, while PPA occlusion produced a slower, progressive decline without reaching ICA levels. Sample size: n = 3–4 per group. Data shown as mean ± SD. Symbols: letters = ICA vs PPA; asterisks (*) = sham vs PPA; exclamation (!) = sham vs ICA. For statistical significance, letters represent ICA vs PPA occlusion, asterisks (*) represents sham vs PPA, exclamation mark (!) represents sham vs ICA; A, ! - P < 0.05, ** - P < 0.01, C, *** - P < 0.001, !!!! – P <0.0001.

### ICA occlusion causes injury to the cortical region

To compare ICA and PPA occlusion in a non-invasive manner, MRI was performed to evaluate rCBF and edema in the cortex and thalamus 24 hours after surgery (Figure 5). Compared to sham (0.93 ± 0.15), rCBF in the cortical region was reduced in both ICA (0.31 ± 0.04) and PPA (0.52 ± 0.058) occluded mice after 15 minutes of ischemia (P < 0.0001) (figure 5A, 5B). When comparing 15 minute-ICA versus 15 minute-PPA occlusions, rCBF was still significantly lower in ICA occluded mice (15 minutes: P < 0.0001). 60 minutes of PPA (0.42 ± 0.086) occlusion did not cause a further drop in rCBF (P = 0.086). Thalamic rCBF showed similar trends, with both ICA (0.38 ± 0.080) and PPA (0.55 ± 0.089) 15-minute occlusion reducing flow compared to sham (0.94 ± 0.045, P <0.0001). Thalamic rCBF was consistently greater in PPA than ICA occlusions at both 15 and 60 minutes (P < 0.05).

**Figure 5.**
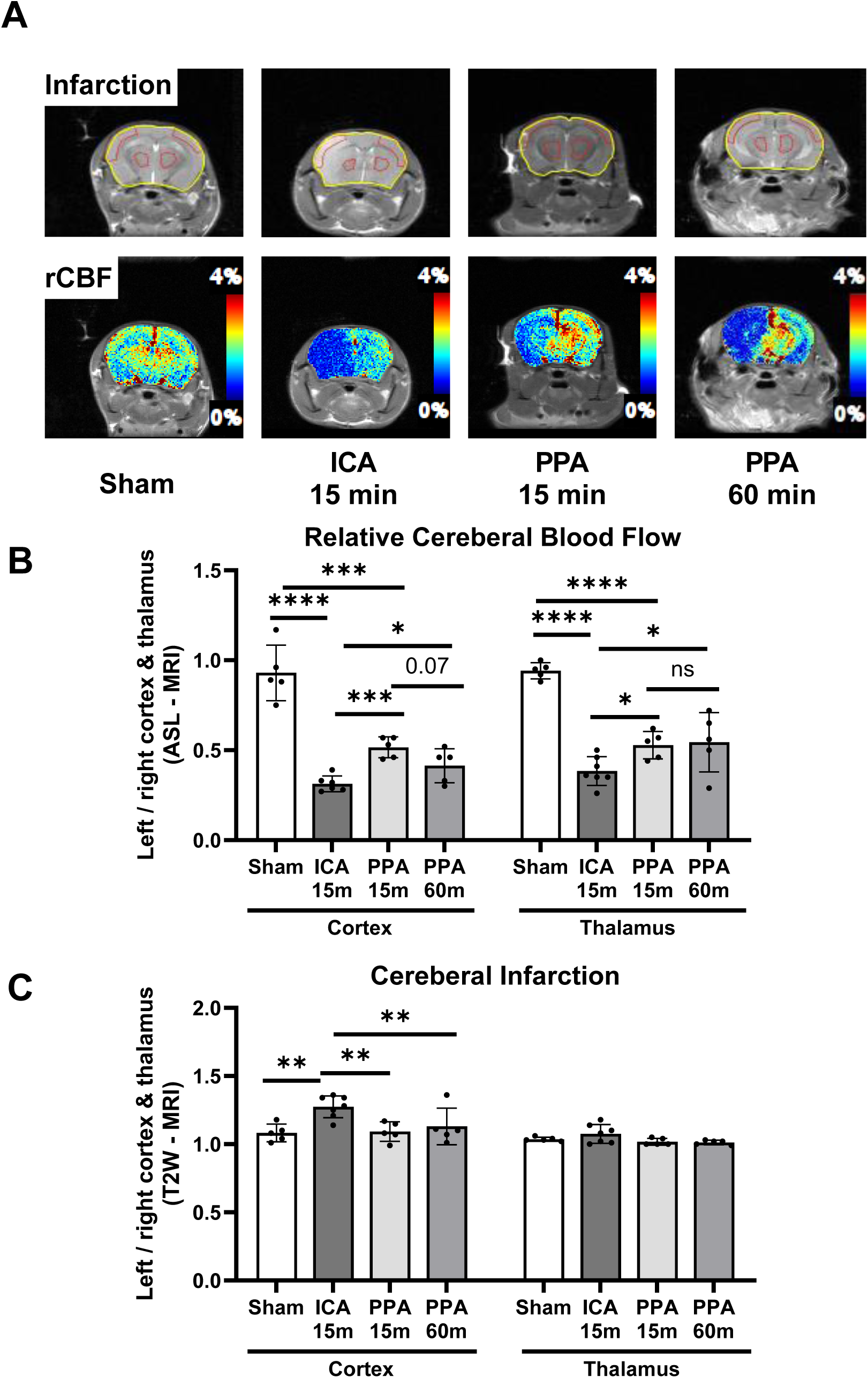
MRI shows ICA occlusion produces cortical edema and greater relative CBF (rCBF) loss. (A) Representative MRI scans of ICA and PPA occluded mice. (B) Quantified rCBF in cortex and thalamus (normalized to contralateral hemisphere). (C) Cortical edema index. ICA occlusion caused marked cortical CBF reduction and edema, while PPA occlusion reduced rCBF without edema. Sample size: n = 5–6 per group. Data shown as mean ± SD. *P < 0.05, **P < 0.01, ***P < 0.001, ****P < 0.0001

When evaluating edema, a predictor of cerebral infarction, 15-minute ICA group developed cortical edema (Sham: 1.08 ± 0.065 vs ICA-15 minutes 1.27 ± 0.080, P = 0.001) (figure 5A, 5C). When compared to sham mice, edema was not observed in PPA occluded mice at 15 minutes (1.093 ± 0.064) or 60 minutes (1.13 ± 0.12) (P > 0.05). Cerebral edema was not detected in the thalamic region for any groups (P > 0.05).

### ICA and PPA occlusion produced a differential inflammatory response

Given the differences in infarct size and BBB permeability, gene expression was assessed for inflammatory cytokines (Figure 6). One-way ANOVA with post-hoc analyses revealed that in IL-6 (sham: 1.054 ± 0.38 vs ICA-30 minutes: 1.67 ± 0.67, P < 0.05, figure 6A) and TGF-β (sham: 1.001 ± 0.061 vs ICA-30 minutes: 1.49 ± 0.54, P < 0.05, figure 6B) were increased after 30 minutes for ICA occlusion. PPA occlusion showed a delayed increase in IL-6 (sham: 0.97 ± 0.096 vs PPA-60 minutes: 1.64 ± 0.70, P < 0.05 figure 6C) with no changes observed for TGF-β levels (figure 6D).

**Figure 6.**
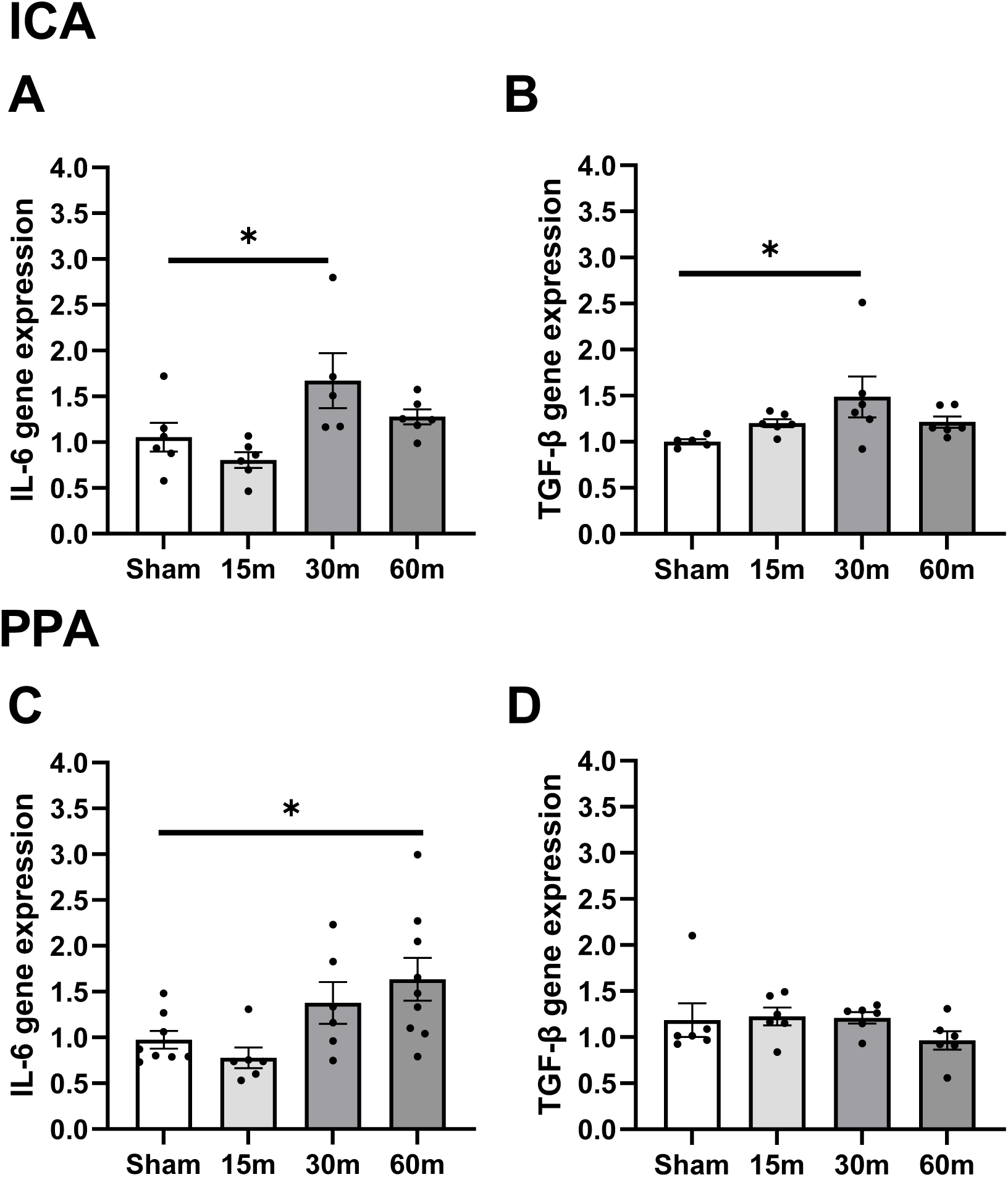
Distinct inflammatory responses after ICA versus PPA occlusion. qPCR quantification of IL-6 and TGF-β. ICA occlusion rapidly upregulated IL-6 and TGF-β, while PPA occlusion increased IL-6 expression at 60 minutes. Sample size: n = 5–9 per group. Data shown as mean ± SD. * - P < 0.05.

## Discussion

This study addressed two overlooked contributors to this variability observed in MCAO studies: (1) the high probability of unintentional filament insertion into the PPA, and (2) the lack of consensus on optimal occlusion duration. By improving the visualization of ICA–PPA bifurcation, this study systematically compared the differences between ICA versus PPA occlusion for the first time. Furthermore, this is the first study to show that the prolonged occlusion of the PPA results in non-specific cerebral injury.

Our findings show that 30 minutes of ICA occlusion represents a critical threshold for generating moderate infarctions (28.11%) while maintaining moderate survival (60% at 24 h). Although longer ICA occlusions (60 min) caused larger infarcts and compromised the BBB, high rates of mortality were observed. Despite the development of milder cerebral injury, improved survivability was observed from 15 minutes ICA occluded mice. In contrast, PPA occlusion resulted in a gradual decline in cerebral blood flow which caused mild cortical injury after 60 minutes of ischemia with high survival rates (10.35% infarct volume, 80% survival). Additionally, the lack of cortical edema, which is commonly observed from MCA occlusions, suggests that the cerebral injury caused from PPA occlusion is non-specific and only occurs after prolonged ischemia.

The modified Koizumi method we developed addresses two major sources of variability: the unintentional occlusion of the PPA and the large range of occlusion times used by previous studies. By mobilizing the hypoglossal nerve, visualization of the ICA–PPA bifurcation is increased resulting in a significant reduction in filament misplacement. Furthermore, our findings reveal that 30 minutes is the optimal occlusion time which balances infarct reproducibility with animal survival. Although it is common to find papers using 60-minute MCAO occlusion ^6,7,30^, the high rates of mortality observed in our study are a significant limitation especially when long term survival is required. Even at 30-minute occlusions, the progressive decline in survival rates after 7 days needs to be taken into consideration by investigators. Although 15 minutes ICA occlusions produce mild cerebral injury, the higher rate of survival may be useful in pharmacological or tissue remodeling studies ^31^.

Our mechanistic analyses also highlight key biological differences. ICA occlusion caused rapid CBF loss, cortical edema, and upregulation of inflammatory cytokines such as IL-6 ^20,32^ and TGF-β ^33,34^. Not only is IL-6 known to be upregulated during thrombosis ^32^ but it has also been reported to be upregulated in serum, cerebrospinal fluid, and stroke tissue in both mice ^35–37^ and humans ^38^. As a pleiotropic cytokine, IL-6 is known to activate the Classical (anti-inflammation) or Trans (pro-inflammation) signaling pathway which influences its functional role in stroke ^36^. While it plays an important role in the recruitment of immune cells during the pro-inflammatory stage ^35,36^, IL-6 has also been reported to be critical during angiogenesis following stroke ^37^. Compared to our study, Gertz *et al* ^37^ reported similar findings where IL-6 gene expression peaked at 2 days following 30 minutes of MCAO occlusion. Additionally, the genetic knockout of IL-6 following MCAO occlusion exacerbated stroke injury due to the impairment of angiogenesis which consequently reduced neovascularization. TGF-β has also been reported in preclinical stroke studies and contributes to the progression of angiogenesis and fibrosis ^33,34^. In contrast, PPA occlusion produced a delayed IL-6 response suggesting a differential inflammatory response which is influenced by the vasculature affected by PPA occlusion. Furthermore, the absence of TGF-β induction in PPA occluded brains further supports that MCA dependent ischemic reperfusion injury engages a broader reparative program which is not triggered by peripheral vascular injury.

Despite the significant reduction in CBF, PPA occlusion did not produce edema in the cortical region suggesting diffuse, non-specific ischemia. These insights not only clarify why prior MCAO studies showed divergent outcomes but also explain why PPA occlusion can be misleadingly interpreted as cerebral infarction. Without the use of imaging techniques such as MRI to confirm cortical injury, researchers may be unaware that they have accidentally occluded the PPA.

This study has several limitations. Firstly, our experiments were conducted in young male C57BL/6 mice. Factors such as age ^39^, sex ^39–42^, and strain ^43^ are known to influence stroke injury^7,44^. For example, premenopausal females show neuroprotective effects which is absent in males ^41^. Secondly, we focused primarily on 24-hour endpoints, which may underestimate longer-term injury or recovery. Finally, while surgical visualization reduces filament misplacement, technical experience remains essential for consistent performance across laboratories. Future studies should extend this work to chronic time points, diverse animal species, and therapeutic intervention testing.

In conclusion, by distinguishing ICA-specific from PPA-driven injury patterns and defining evidence-based occlusion parameters, this work provides a framework for future MCAO studies. The marked differences observed between acute (15 minutes) and prolonged (30–60 minutes) occlusion underscore the importance of careful study design and verification that the MCAO model has been performed correctly. By identifying two critical sources of variability in the MCAO model we hope this study provides clear, actionable guidance for reproducible and translationally relevant stroke research.

## Non-standard Abbreviations and Acronyms

BBB: blood-brain barrier
CBF: Cerebral Blood Flow
CCA: Common carotid artery
HG: Hypoglossal
ICA: Internal carotid artery
IL-6: Interleukin 6
LCCA: Left common artery
LECA: Left external carotid artery
LICA: Left internal carotid artery
MCA: Middle cerebral artery
MCAO: middle cerebral artery occlusion
MRI: Magnetic Resonance Imaging
PPA: pterygopalatine artery
rCBF: relative cerebral blood flow
TGF-β: Tumor Growth Factor Beta
TTC: 2,3,5-Triphenyltetrazolium chloride

