## Supplemental file for "Standardizing the mouse MCAO Stroke Model: Distinguishing Internal Carotid vs Pterygopalatine Artery Occlusion"

Supplemental material

Methods

Middle Cerebral Artery Occlusion (MCAO) mouse model

**Reagents / Consumables**

- Surgical tools
  - Graefe Forceps (RS-5138)
  - Student Vannas Spring Scissors (91500-09)
  - Delicate Suture Tying Forceps (11063-07)
  - Castroviejo Micro Needle Holder (6006-50)
  - Symmetry Scissors Westcott Tenotomy (54-6513)
- Intubation catheter – 20G IV catheter (Exel International, Virginia, USA)
- Fine MCAO suture monofilaments coated in white silicone rubber – (Doccol Corporation, Massachusetts, USA)
  - 7019910PK5Re – Diameter: 0.19 mm, Silicone coating length: 9-10 mm
  - 6021910PK5Re – Diameter: 0.21 mm, Silicone coating length: 9-10 mm
- Sutures
  - 7-0 polypropylene (Sharpoint)
  - 5-0 silk (COVIDIEN)
- Rodent hair shaver
- Ethanol 70%

**Equipment**

- SM-3T series Zoom Stero Microscope - (Amscope, California, USA)
- PN 118918 Rev 04 XGI-8 Veterinary Gas Anesthesia System (Caliper Life Sciences, Massachusetts, USA)
- VIP 3000 Veterinary Vaporizer (Midmark, Ohio, USA)
- Induction chamber
- Heated Surgical Table - (Harvard Apparatus, Massachusetts, USA)
- Heat lamp
- Medical grade oxygen
- Rodent intubation stand
- Minivent Ventilator (Model 845) – (Harvard Apparatus, Massachusetts, USA)

**Procedure**

1. Mice are weighed, and placed into an isoflurane induction chamber which is filled with 3% isoflurane using a VIP3000 vaporizer in conjunction with a XGI-8 Veterinary Gas Anesthesia system, until complete anesthesia is achieved.
2. Transfer the anesthetized animal onto a rodent intubation stand and intubate the animal with a 20G angiocath sleeve with the assistance of a SM-3T series Zoom Stero Microscope.
3. The mouse is placed onto a preheated Harvard Apparatus heated surgical table at 37°C, supine position, and connected to a Minivent Ventilator (Model 845). Tidal volume (Vt; mL) is maintained at 148 mL, whereas respiratory rate ($min^{-1}$) is maintained at 150 bpm. Surgical anesthesia (isoflurane, 1.0% at a flow rate of 0.5-1.0 L/min) is maintained through the same intubation line.
4. Once complete anesthesia is confirmed through paw reflex pinch test, the neck area is shaved and aseptically prepared with betadine solution with 70% isopropyl solution.
5. Following a midline neck incision, a blunt dissection is performed with Graefe forceps to pull apart the soft tissue until the left common artery (LCCA) is identified.
6. The nerves surrounding the LCCA are then carefully dissected through blunt dissection, and a ligature is made with 7-0 polypropylene with delicate suture tying forceps.
7. Identify the left external carotid artery (LECA) and the left internal carotid artery (LICA) was carefully separate from the surrounding tissue.
8. To maximize the visibility of the LICA and the left pterygopalatine artery (LPPA), the hypoglossal (HG) nerve is mobilized by carefully dissecting the surrounding connective tissue around the HG nerve (figure S1). Do not directly touch the HG nerve.
9. The LECA is temporarily occluded with a reversible 5.0 silk ligature whereas a loose 5.0 silk ligature is prepared around the LICA using delicate suture tying forceps.

**Formatted:** Font: (Default) Times New Roman, Not Bold, Not Italic

10. Before the LCCA bifurcates to the LECA and the LICA, a small incision is made into the LCCA using a pair of symmetry scissors.
11. Depending on the size of the animal, a Silicone Rubber Coated monofilament (for 15–20-gram mice, filament diameter: 0.19 mm. For 21-25 g mice, 6021910PK5Re – filament diameter: 0.21 mm) is inserted into the incision. The monofilament is threaded through the LCCA until it reaches the branching point between the LICA and the left pterygopalatine artery (LPPA).
12. For LCCA occlusion, the filament is manipulated (see supplemental video) to ensure that it enters the LICA. Continue inserting the filament until the first sensation of resistance is detected and secure the filament with the previously prepared ligature prepared on the LICA[A].
    A. The depth of the filament is critical to ensure that consistent cerebral infarction. Going too deep can lead to mortality while a shallow insertion will reduce the severity of cerebral ischemia.
    B. Place a drop of sterile PBS on the surgical site to prevent the tissue from drying out.
13. After the occlusion is maintained for the required amount of time, the LCCA ligature is loosened, and the filament is withdrawn to restore blood flow into the occluded region. The LCCA is then permanently tied off with 5.0 silk suture.
14. After the muscular tissue and skin is closed with 5.0 silk suture, the mouse is weaned from isoflurane. Once voluntary respiration is observed, the animal is removed from the ventilator, placed under a heatlamp, and supplemented with medical grade oxygen to assist in recovery.

15. For neurological deficiency, TTC staining, assessment of blood brain barrier function, or molecular analyses, brains can be collected after 24 hours.

Table S1

| **Study Title** | **DOI** | **Occlusion Time** |
|---|---|---|
| Melatonin enhances neurogenesis and neuroplasticity in long-term recovery following cerebral ischemia in mice | 10.1016/j.bbadis.2025.167738. | 30 min |
| Shengui Sansheng San alleviates the worsening of blood-brain barrier integrity resulted from delayed tPA administration through VIP/VIPR1 pathway | 10.1186/s13020-025-01079-0. | 300 min |
| Periorbital Placement of a Laser Doppler Probe for Cerebral Blood Flow Monitoring Prior to Middle Cerebral Artery Occlusion in Rodent Models | 10.3791/66839. | 60 min |
| Biological and Procedural Predictors of Outcome in the Stroke Preclinical Assessment Network (SPAN) Trial | 10.1161/CIRCRESAHA.123.324139. | 60 min |
| High-resolution micro-CT for 3D infarct characterization and segmentation in mice stroke models | 10.1038/s41598-022-21494-9. | 10 min and 45 min |
| N-Methyl-D-Aspartate Receptors Antagonist Prevents Secondary Ischemic Brain Injury Associated With Lipopolysaccharide-Induced Sepsis-Like State Presumably via Immunomodulatory Actions | 10.3389/fncel.2022.881088. | 30 min |
| Transient Middle Cerebral Artery Occlusion with an Intraluminal Suture Enables Reproducible Induction of Ischemic Stroke in Mice | 10.21769/BioProtoc.4305. | 30, 45, 60, or 90 min |
| Post-ischemic protein restriction induces sustained neuroprotection, neurological recovery, brain remodeling, and gut microbiota rebalancing | 10.1016/j.bbi.2021.11.016 | 30 min |
| Neuroprotective effects of minocycline and KML29, a potent inhibitor of monoacylglycerol lipase, in an experimental stroke model: a small-animal positron emission tomography study | 10.7150/thno.64320 | 30 min |
| Small extracellular vesicles obtained from hypoxic mesenchymal stromal cells have unique characteristics that promote cerebral | 10.1007/s00395-021-00881-9. | 40 min |

| | | |
|---|---|---|
| angiogenesis, brain remodeling and neurological recovery after focal cerebral ischemia in mice | | |
| Hypocaloric Diet Initiated Post-Ischemia Provides Long-Term Neuroprotection and Promotes Peri-Infarct Brain Remodeling by Regulating Metabolic and Survival-Promoting Proteins | 10.1007/s12035-020-02207-7. | 30 min |
| Light Sheet Microscopy Using FITC-Albumin Followed by Immunohistochemistry of the Same Rehydrated Brains Reveals Ischemic Brain Injury and Early Microvascular Remodeling | 10.3389/fncel.2020.625513. | 30 min |
| Stroke Severity, and Not Cerebral Infarct Location, Increases the Risk of Infection | 10.1007/s12975-019-00738-3. | check |
| N,N-dimethyltryptamine reduces infarct size and improves functional recovery following transient focal brain ischemia in rats | 10.1016/j.expneurol.2020.113245. | 60 min |
| Systemic conditioned medium treatment from interleukin-1 primed mesenchymal stem cells promotes recovery after stroke | 10.1186/s13287-020-1560-y. | 15 min, 20 min |
| Moderate Protein Restriction Protects Against Focal Cerebral Ischemia in Mice by Mechanisms Involving Anti-inflammatory and Anti-oxidant Responses | 10.1007/s12035-019-01679-6. | 30 min |
| Proteomic Analysis of Rat Cerebral Cortex in the Subacute to Long-Term Phases of Focal Cerebral Ischemia-Reperfusion Injury | 10.1021/acs.jproteome.9b00220. | 120 min |
| Behavioral tests that reveal long-term deficits after permanent focal cerebral ischemia in mouse | 10.1016/j.bbr.2018.11.040. | Permanent MCAO |
| A heparan sulfate-based matrix therapy reduces brain damage and enhances functional recovery following stroke | 10.7150/thno.28252 | 60 min |
| Metabolomic Analysis of Mouse Brain after a Transient Middle Cerebral Artery Occlusion by Mass Spectrometry Imaging | 10.2176/nmc.oa.2018-0054. | 60 min |
| Effect of laser Doppler flowmetry and occlusion time on outcome variability and mortality in rat middle cerebral artery occlusion: inconclusive results | 10.1186/s12868-018-0425-0. | 45 min |
| Oral administration of a novel lipophilic PPARδ agonist is not neuroprotective after rodent cerebral ischemia | 10.1177/0271678X17743876. | 45 min |
| Multi-site laser Doppler flowmetry for assessing collateral flow in experimental ischemic stroke: Validation of outcome prediction with acute MRI | 10.1177/0271678X16661567. | 90 min |
| A specific dietary intervention to restore brain structure and function after ischemic stroke | 10.7150/thno.17559 | 30 min |

| | | |
|---|---|---|
| Investigating potentially salvageable penumbra tissue in an in vivo model of transient ischemic stroke using sodium, diffusion, and perfusion magnetic resonance imaging | 10.1186/s12868-016-0316-1. | 90 min |
| Evaluation of a filament perforation model for mouse subarachnoid hemorrhage using 7.0 Tesla MRI | 10.1016/j.jocn.2015.10.045. | 30 min |
| Systemic inflammation affects reperfusion following transient cerebral ischaemia | 10.1016/j.expneurol.2016.01.013. | 30 min |
| A Comparative Study of Variables Influencing Ischemic Injury in the Longa and Koizumi Methods of Intraluminal Filament Middle Cerebral Artery Occlusion in Mice | 10.1371/journal.pone.0148503. | 30 min |
| Quantitative T2* mapping reveals early temporo-spatial dynamics in an ischemic stroke model | 10.1016/j.jneumeth.2015.11.018. | 60min |
| Ultrasonic vocalization in murine experimental stroke: A mechanistic model of aphasia | 10.3233/RNN-150583. | 45 min |
| Delayed reperfusion deficits after experimental stroke account for increased pathophysiology | 10.1038/jcbfm.2014.197. | 30 min |
| Preclinical evaluation of recombinant T cell receptor ligand RTL1000 as a therapeutic agent in ischemic stroke | 10.1007/s12975-014-0373-7. | 60 min |
| Isoflurane reduces the ischemia reperfusion injury surge: a longitudinal study with MRI | 10.1016/j.brainres.2014.08.003 | 90 min |
| Severity of middle cerebral artery occlusion determines retinal deficits in rats | 10.1016/j.expneurol.2014.02.005. | 90 min |
| Keep warm and get success: the role of postischemic temperature in the mouse middle cerebral artery occlusion model | 10.1016/j.brainresbull.2013.12.003. | 90 min |
| Spatiotemporal uptake characteristics of [18]F-2-fluoro-2-deoxy-D-glucose in a rat middle cerebral artery occlusion model | 10.1161/STROKEAHA.113.000903 | 150 min |
| Anticoagulation with dabigatran does not increase secondary intracerebral haemorrhage after thrombolysis in experimental cerebral ischaemia | 10.1160/TH12-12-0942. | 120 and 180 min |
| Mild hypothermia reduces tissue plasminogen activator-related hemorrhage and blood brain barrier disruption after experimental stroke | 10.1089/ther.2013.0010. | 24 hours |
| Brain infarct volume after permanent focal ischemia is not dependent on Nox2 expression | 10.1016/j.brainres.2012.09.023. | 24 hr |
| Consistent focal cerebral ischemia without posterior cerebral artery occlusion and its real-time monitoring in an intraluminal suture model in mice | 10.3171/2011.11.JNS111167. | 60 min |

| | | |
|---|---|---|
| Intraluminal middle cerebral artery occlusion (MCAO) model for ischemic stroke with laser doppler flowmetry guidance in mice | 10.3791/2879. | 45 min |
| Increased Risk of Hemorrhagic Transformation in Ischemic Stroke Occurring During Warfarin Anticoagulation: An Experimental Study in Mice | 10.1161/STROKEAHA.110.60 4652. | 180 min |
| Combined contrast-enhanced ultrasound and rt-PA treatment is safe and improves impaired microcirculation after reperfusion of middle cerebral artery occlusion | 10.1038/jcbfm.2010.82. | 90 min |
| Human microglia transplanted in rat focal ischemia brain induce neuroprotection and behavioral improvement | 10.1371/journal.pone.0011746. | 90 min |
| Pre-ischemic exercise preserves cerebral blood flow during reperfusion in stroke | 10.1179/016164109X1258109 6796431. | 30 minutes |
| Microvessel changes after post-ischemic benign and malignant hyperemia: | 10.1186/1471-2377-10-24. | 30 min, 180 min, 360 min |
| An optimized mouse model for transient ischemic attack. | 10.1097/NEN.0b013e3181cd3 31c. | 20 min |
| Neither in vivo MRI nor behavioural assessment indicate therapeutic efficacy for | 10.1186/1471-2202-10-82. | 60 min |
| Microvessel changes after post-ischemic benign and malignant hyperemia: experimental study in rats | 10.1097/ALN.0b013e3181a1fe 68. | 45 min, 60 min |
| Ginkgo biloba extract neuroprotective action is dependent on heme oxygenase 1 in ischemic reperfusion brain injury | 10.1161/STROKEAHA.108.52 3480. | 90 min, 120 min |
| LAU-0901, a novel platelet-activating factor antagonist, is highly neuroprotective in cerebral ischemia | 10.1016/j.expneurol.2008.08.0 09. | 120 min |
| Hyperthermia induced after recirculation triggers chronic neurodegeneration in the penumbra zone of focal ischemia in the rat brain | 10.1590/s0100-879x2008001100014. | 60 min |
| Delayed tolerance with repetitive transient focal ischemic preconditioning in the mouse | 10.1161/STROKEAHA.107.49 7412 | 45 min, 90 min |
| Platelet adhesion receptors do not modulate infarct volume after a photochemically induced stroke in mice | 10.1016/j.brainres.2007.07.103 . | 90 min |
| Delayed hyperbaric oxygenation is more effective than early prolonged normobaric hyperoxia in experimental focal cerebral ischemia | 10.1016/j.neulet.2007.07.009. | 150 min |
| Infarct volume after transient middle cerebral artery occlusion (MCAo) can be reduced by attenuation but not by inactivation of c-Jun action | 10.1016/j.brainres.2007.03.023 . | 90 min |

| | | |
|---|---|---|
| Therapeutic time window of neuroprotection by non-competitive AMPA antagonists in transient and permanent focal cerebral ischemia in rats | 10.1016/j.brainres.2006.09.043. | Permanent MCAO |
| Intravenous administration of glial cell line-derived neurotrophic factor gene-modified human mesenchymal stem cells protects against injury in a cerebral ischemia model in the adult rat | 10.1002/jnr.21056. | Permanent MCAO |
| Tacrolimus (FK506) attenuates biphasic cytochrome c release and Bad phosphorylation following transient cerebral ischemia in mice | 10.1016/j.neuroscience.2006.06.064. | 60 min |
| Therapeutic window of bradykinin B2 receptor inhibition after focal cerebral ischemia in rats | 10.1016/j.neuint.2006.02.010. | 90 min |
| Oxygen therapy in permanent brain ischemia: potential and limitations | 10.1016/j.brainres.2006.05.108. | Permanent MCAO |
| Modest MRI signal intensity changes precede delayed cortical necrosis after transient focal ischemia in the rat | 10.1161/01.STR.0000221713.06148.16. | 60 min |
| Resveratrol reduces the elevated level of MMP-9 induced by cerebral ischemia-reperfusion in mice | 10.1016/j.lfs.2005.10.030. | 60 min |
| Allopregnanolone, a progesterone metabolite, is more effective than progesterone | 10.1016/j.annemergmed.2005.12.011. | 120 min |
| Hyperbaric oxygen reduces basal lamina degradation after transient focal cerebral ischemia in rats | 10.1016/j.brainres.2006.01.013. | 45 min |
| Dietary phytoestrogens improve stroke outcome after transient focal cerebral ischemia in rats | 10.1111/j.1460-9568.2006.04599.x. | 90 min |
| Alpha-linolenic acid and riluzole treatment confer cerebral protection and improve survival after focal brain ischemia | 10.1016/j.neuroscience.2005.08.083. | 60 mins |
| Temporal profile of T2-weighted MRI distinguishes between pannecrosis and selective neuronal death after transient focal cerebral ischemia in the rat | 10.1038/sj.jcbfm.9600166. | 60 mins |
| Social interaction improves experimental stroke outcome | 10.1161/01.STR.0000177538.17687.54. | 60 mins or 90 mins |
| Hyperbaric oxygen reduces blood-brain barrier damage and edema after transient focal cerebral ischemia | 10.1161/01.STR.0000173408.94728.79. | 120 min |
| Aggravation of focal cerebral ischemia by tissue plasminogen activator is reversed by 3-hydroxy-3-methylglutaryl coenzyme A reductase inhibitor but does not depend on endothelial NO synthase | 10.1161/01.STR.0000152273.24063.f7. | 90 min |
| Post-ischemic delivery of the 3-hydroxy-3-methylglutaryl coenzyme A reductase inhibitor | 10.1016/j.neuroscience.2005.04.063. | 90 min |

| | | |
|---|---|---|
| rosuvastatin protects against focal cerebral ischemia in mice via inhibition of extracellular-regulated kinase-1/-2 | | |
| The intraluminal thread model revisited: rat strain differences in local cerebral blood flow | 10.1179/016164105X18214. | 60 min |
| Characterizing the diffusion/perfusion mismatch in experimental focal cerebral ischemia | 10.1002/ana.10803. | 60 min or permanent |
| Neuroprotective effects of insulin-like growth factor-binding protein ligand inhibitors in vitro and in vivo | 10.1097/01.WCB.0000087091.01171.AE. | permanent |
| Combination drug therapy and mild hypothermia after transient focal cerebral ischemia in rats | 10.1161/01.STR.0000083622.65684.21. | 90 min |
| Closure of the blood-brain barrier by matrix metalloproteinase inhibition reduces rtPA-mediated mortality in cerebral ischemia with delayed reperfusion | 10.1161/01.STR.0000083051.93319.28. | 90 min, 180 min, 300 min |
| Characterization of a new double-filament model of focal cerebral ischemia in heme oxygenase-2-deficient mice | 10.1152/ajpregu.00067.2003. | 60 mins or 120 mins |
| Neuroprotective effect of SolCD39, a novel platelet aggregation inhibitor, on transient middle cerebral artery occlusion in rats | 10.1161/01.STR.0000056169.45365.15. | 120 mins |
| Intravenous TAT-Bcl-Xl is protective after middle cerebral artery occlusion in mice | 10.1002/ana.10356. | 30 mins, 90 min |
| Neuroprotective effect of delayed moderate hypothermia after focal cerebral ischemia: an MRI study | 10.1161/01.str.0000019603.29818.9c. | 120 min |
| Social stress exacerbates focal cerebral ischemia in mice | 10.1161/01.str.0000016967.76805.bf. | 60 mins |
| Normobaric hyperoxia reduces MRI diffusion abnormalities and infarct size in experimental stroke | 10.1212/wnl.58.6.945. | 120 min |
| Filament size influences temperature changes and brain damage following middle cerebral artery occlusion in rats | 10.1007/s00221-001-0909-4. | 120 min |
| Phosphorylation of cyclic adenosine monophosphate response element binding protein in oligodendrocytes in the corpus callosum after focal cerebral ischemia in the rat | 10.1097/00004647-200110000-00006. | 90 min |
| Dehydroascorbic acid, a blood-brain barrier transportable form of vitamin C, mediates potent cerebroprotection in experimental stroke | 10.1073/pnas.171325998. | 45 min or permanent |

| | | |
|---|---|---|
| Anesthetic choice of halothane versus propofol: impact on experimental perioperative stroke | 10.1161/01.str.32.8.1920. | 120 min |
| Intranasal administration of insulin-like growth factor-I bypasses the blood-brain barrier and protects against focal cerebral ischemic damage | 10.1016/s0022-510x(01)00532-9. | 120 min |
| Mevastatin, an HMG-CoA reductase inhibitor, reduces stroke damage and upregulates endothelial nitric oxide synthase in mice | 10.1161/01.str.32.4.980. | 120 min |
| Neuroprotection by intrathecal application of liposome-entrapped fasudil in a rat model of ischemia | 10.2176/nmc.41.107. | 120 min |
| Cognitive deficits after focal cerebral ischemia in mice. | 10.1161/01.str.31.8.1939. | 60 min, 90 min |
| Lack of interleukin-6 expression is not protective against focal central nervous system ischemia | 10.1161/01.str.31.7.1715. | 45 min, 120 min |
| Hypertonic saline worsens infarct volume after transient focal ischemia in rats | 10.1161/01.str.31.7.1694. | 120 min |
| Delayed systemic administration of PACAP38 is neuroprotective in transient middle cerebral artery occlusion in the rat | 10.1161/01.str.31.6.1411. | 120 min |
| Persistent neuroprotection with prolonged postischemic hypothermia in adult rats subjected to transient middle cerebral artery occlusion | 10.1006/exnr.2000.7369. | 120 min |
| Immunohistochemical detection of leukemia inhibitory factor after focal cerebral ischemia in rats | 10.1097/00004647-200004000-00003. | 90 min |
| sigma(1)-receptor ligand 4-phenyl-1-(4-phenylbutyl)-piperidine affords neuroprotection from focal ischemia with prolonged reperfusion | 10.1161/01.str.31.4.976. | 120 min |
| Increased expression of intercellular adhesion molecule-1 in mouse focal cerebral ischemia model | Not found | Permanent MCAO |
| Spontaneous hyperthermia and its mechanism in the intraluminal suture middle cerebral artery occlusion model of rats | 10.1161/01.str.30.11.2464. | 60 min, 90 min, 120 min, permanent |
| Titration of postischemic cerebral hypoperfusion by variation of ischemic severity in a murine model of stroke | 10.1097/00006123-199908000-00027. | 15 min, 30 min, 45 min |
| Middle cerebral artery occlusion in the mouse by intraluminal suture coated with poly-L-lysine: neurological and histological validation | 10.1016/s0006-8993(99)01528-0. | 30 min, 60 min, 120 min, 180 min |

| | | |
|---|---|---|
| Time course of IL-6 expression in experimental CNS ischemia | 10.1080/01616412.1999.11740933. | 120 min |
| Effects of postischemic halothane administration on outcome from transient focal cerebral ischemia in the rat | 10.1097/00008506-199901000-00006. | 75 minutes |
| Neuroprotection from focal ischemia by 4-phenyl-1-(4-phenylbutyl) piperidine (PPBP) is dependent on treatment duration in rats | 10.1097/00000539-199812000-00016. | 120 min |
| Monofilament intraluminal middle cerebral artery occlusion in the mouse | 10.1080/01616412.1997.11740874. | 60 min, 120 min, 180 min or permanent |
| Extracellular potassium in a neocortical core area after transient focal ischemia | 10.1161/01.str.28.1.206. | 120 min |
| Neuroprotective effects of preischemia intraarterial magnesium sulfate in reversible focal cerebral ischemia | 10.3171/jns.1996.85.1.0117. | 90 min, 120 min |
| A comparison of the early development of ischemic brain damage in normoglycemic and hyperglycemic rats using magnetic resonance imaging | 10.1007/BF00228624. | 60 min or permanent |
| Graded hypotension and MCA occlusion duration: effect in transient focal ischemia | 10.1038/jcbfm.1995.124. | 120 min |
| Delayed treatment with intravenous basic fibroblast growth factor reduces infarct size following permanent focal cerebral ischemia in rats | 10.1038/jcbfm.1995.121. | permanent |
| N-tert-butyl-alpha-phenylnitrone improves recovery of brain energy state in rats following transient focal ischemia | 10.1073/pnas.92.11.5057. | 120 min |
| Combined perfusion and diffusion-weighted magnetic resonance imaging in a rat model of reversible middle cerebral artery occlusion | 10.1161/01.str.26.3.451. | 45 min, 120 min |
| Temporal evolution and spatial distribution of the diffusion constant of water in rat brain after transient middle cerebral artery occlusion | 10.1016/0022-510x(93)90262-w. | 120 min |
| Postischemic (1 hour) hypothermia significantly reduces ischemic cell damage in rats subjected to 2 hours of middle cerebral artery occlusion | 10.1161/01.str.24.8.1235. | 120 min |
